# Bacterial motility depends on a critical flagellum length and energy-optimised assembly

**DOI:** 10.1101/2024.06.28.599820

**Authors:** Manuel Halte, Philipp F. Popp, David Hathcock, John Severn, Svenja Fischer, Christian Goosmann, Adrien Ducret, Emmanuelle Charpentier, Yuhai Tu, Eric Lauga, Marc Erhardt, Thibaud T. Renault

## Abstract

The flagellum is the most complex macromolecular structure known in bacteria and comprised of around two dozen distinct proteins. The main building block of the long, external flagellar filament, flagellin, is secreted through the flagellar type-III secretion system at a remarkable rate of several tens of thousands amino acids per second, significantly surpassing the rates achieved by other pore-based protein secretion systems. The evolutionary implications and potential benefits of this high secretion rate for flagellum assembly and function, however, have remained elusive. In this study, we provide both experimental and theoretical evidence that the flagellar secretion rate has been evolutionarily optimized to facilitate rapid and efficient construction of a functional flagellum. By synchronizing flagellar assembly, we found that a minimal filament length of 2.5 µm was required for swimming motility. Biophysical modelling revealed that this minimal filament length threshold resulted from an elasto-hydrodynamic instability of the whole swimming cell, dependent on the filament length. Furthermore, we developed a stepwise filament labeling method combined with electron microscopy visualization to validate predicted flagellin secretion rates of up to 10,000 amino acids per second. A biophysical model of flagellum growth demonstrates that the observed high flagellin secretion rate efficiently balances filament elongation and energy consumption, thereby enabling motility in the shortest amount of time. Taken together, these insights underscore the evolutionary pressures that have shaped the development and optimization of the flagellum and type-III secretion system, illuminating the intricate interplay between functionality and efficiency in assembly of large macromolecular structures.

**Significance statement:** Our study demonstrates how protein secretion of the bacterial flagellum is finely tuned to optimize filament assembly rate and flagellum function while minimizing energy consumption. By measuring flagellar filament lengths and bacterial swimming after initiation of flagellum assembly, we were able to establish the minimal filament length necessary for swimming motility, which we rationalized physically as resulting from an elasto-hydrodynamic instability of the swimming cell. Our biophysical model of flagellum growth further illustrates how the physiological flagellin secretion rate is optimized to maximize filament elongation while conserving energy. These findings illuminate the evolutionary pressures that have shaped the function of the bacterial flagellum and type-III secretion system, driving improvements in bacterial motility and overall fitness.

## Introduction

Most bacteria use the flagellum, the largest self-assembling protein structure known in bacteria, as their primary motility device (1). It is produced in consecutive steps that are regulated on multiple levels (2, 3). The hook-basal body (HBB) that contains the flagellar secretion system has a molecular weight of ∼ 15 MDa in *Salmonella enterica* (4). After assembly of the rod that spans the bacterial cell envelope, the hook is assembled as the first extra-cellular structure protruding about 55 nm outside of the cell (5). Subsequently, the long filament with a molecular weight of ∼ 110 MDa per µm self-assembles from a single protein, flagellin. The hook and flagellum, helical and hollow structures, are distinct in their mechanical properties: the hook is supercoiled and flexible, while the flagellum is supercoiled and rigid (6). The rotation of the flagellar motor consisting of the membrane proteins MotAB (stator) and FliG (rotor) in *S. enterica* (7) is transmitted to the rigid filament via the flexible hook that acts as a universal joint. The physics underlying this movement involves the rotation of each motor inducing rotation of the hook and the flagellar filament attached to it, thereby generating propulsion in viscous fluids (1, 8).

While assembly of the bacterial flagellum is well understood from a structural perspective, many questions re-main concerning the molecular mechanisms of protein secretion. Most components of the flagellum, including the rod, the hook and the filament subunits, are substrates of the flagellum-specific type-III secretion system (fT3SS) that is located at the base of the HBB structure (9, 10). Translocation of flagellar substrates across the cytoplasmic membrane depends on the electrochemical gradient of protons (proton motive force, pmf) and is likely mediated via the basal body membrane protein FlhA (11, 12). The external flagellar filament can grow to a length of 10-20 µm, corresponding to ∼ 20,000 to 40,000 flagellin subunits (13). Several models have been proposed to explain the energization of flagellin export and assembly of the filament subunits outside of the cell (11, 14–17). Although the exact mechanism underlying energetics of protein translocation remains unclear, the pmf-dependent pumping activity of the fT3SS results in an injection force, which propels the substrates across the cytoplasmic membrane into the 2 nm wide secretion channel of the flagellum, where they further diffuse to reach their site of assembly at the distal end of the growing structure (13, 14, 18, 19).

Assembly of the bacterial filament itself is surprisingly fast; in a previous study, we used real-time fluorescence microscopy and stepwise fragment labeling to reveal the kinetics of flagellar filament elongation (18). We demonstrated that the filament growth rate is initially very fast (∼100 nm/min, corresponding to 3.5 flagellins/s or ∼1,700 amino acids per second) and quickly decreases in a non-linear way to fall below 20 nm/min. Interestingly, however, the theoretical injection rate of the first flagellin molecule, the *k*_on_ parameter in the kinetic injection diffusion model (18), was estimated to be 30 s^-1^ (or 30 flagellins/s, equivalent to ∼15,000 amino acids per second). Experimental determination of the *k*_on_ parameter is not directly possible by measuring filament elongation using the stepwise labeling approach, since what is measured is an average speed based on increments of the filament length over a defined time, which inevitably underestimates the true parameter. We note that the theoretical secretion rate of the fT3SS is several orders of magnitude higher than that of other protein secretion pores (*e.g.* 16 amino acids per second in the type-I secretion system (20), 40 amino acids per second in the Sec translocon (21)), raising the question of the biological relevance of such a high secretion rate. The ability to move is of paramount importance for bacteria, as motility provides substantial fitness benefits such as enhanced nutrient acquisition, avoidance of toxic substances and effective colonization of hosts and surfaces (22–25). It therefore appears reasonable to speculate that enabling fast assembly of the flagellar filament while balancing the associated energy costs of a rapid secretion process would appear beneficial in order to acquire the ability to move as soon as possible.

Here, we combine analyses of single cell swimming velocities and flagellar filament lengths to reveal a swift transition between non-swimming and swimming bacterial sub-populations at a filament length threshold of ∼ 2.5 µm. We apply these findings to a biophysical model of swimming bacteria to rationalize how a minimal filament length of ∼ 2.5 µm is sufficient to enable swimming motility. We further develop an improved filament labeling method using electron microscopy and short elongation time (down to 30 s) to experimentally validate the predicted rapid flagellin secretion rate of ∼ 10,000 amino acids per second. In order to rationalize the costs/benefits of a such rapid flagellin secretion rate, we develop a biophysical model of flagellum growth that describes the energy costs associated with flagellin secretion with measurements of swimming motility and reveals that the observed rapid flagellin secretion rate is within the range of optimal efficiency.

## Results

### Minimal flagellar filament length required for motility

Unlike the flagellar rod, hook, and injectisome needle, there is no direct mechanism controlling filament length (26). It is commonly accepted that while flagellar filaments grow up to 15-20 µm, a much shorter length (at least one helical pitch, or about 2 µm) is sufficient for bacteria to swim and to allow changing direction (27–29). Thus, it appeared reasonable to assume that long filaments are merely a byproduct of the growth kinetics and that the delay in becoming motile is rather the biologically important parameter. To address this question experimentally, we used an anhydrotetracycline (AnTc) inducible flagellar master regulator (P*tetA-flhDC*) to synchronize flagella biosynthesis (18). This allowed us to simultaneously track flagellar filament growth and observe the swimming behavior of individual bacteria (18) (Figure 1A). After induction of flagellar gene expression using addition of AnTc, we recorded the fraction of motile bacteria and their swimming speed while determining the lengths of flagellar filaments using immunostaining (Figure 1A). We observed that virtually all bacteria within the population (>90%) were flagellated after 30 min post-induction. Swimming motility was observed after 40 min post induction, and swimming speed sharply increased between 50 and 60 min post induction before reaching a maximal speed of ∼ 30 µm/s after 80 min post induction (Figure 1B and C).

**Figure 1.**
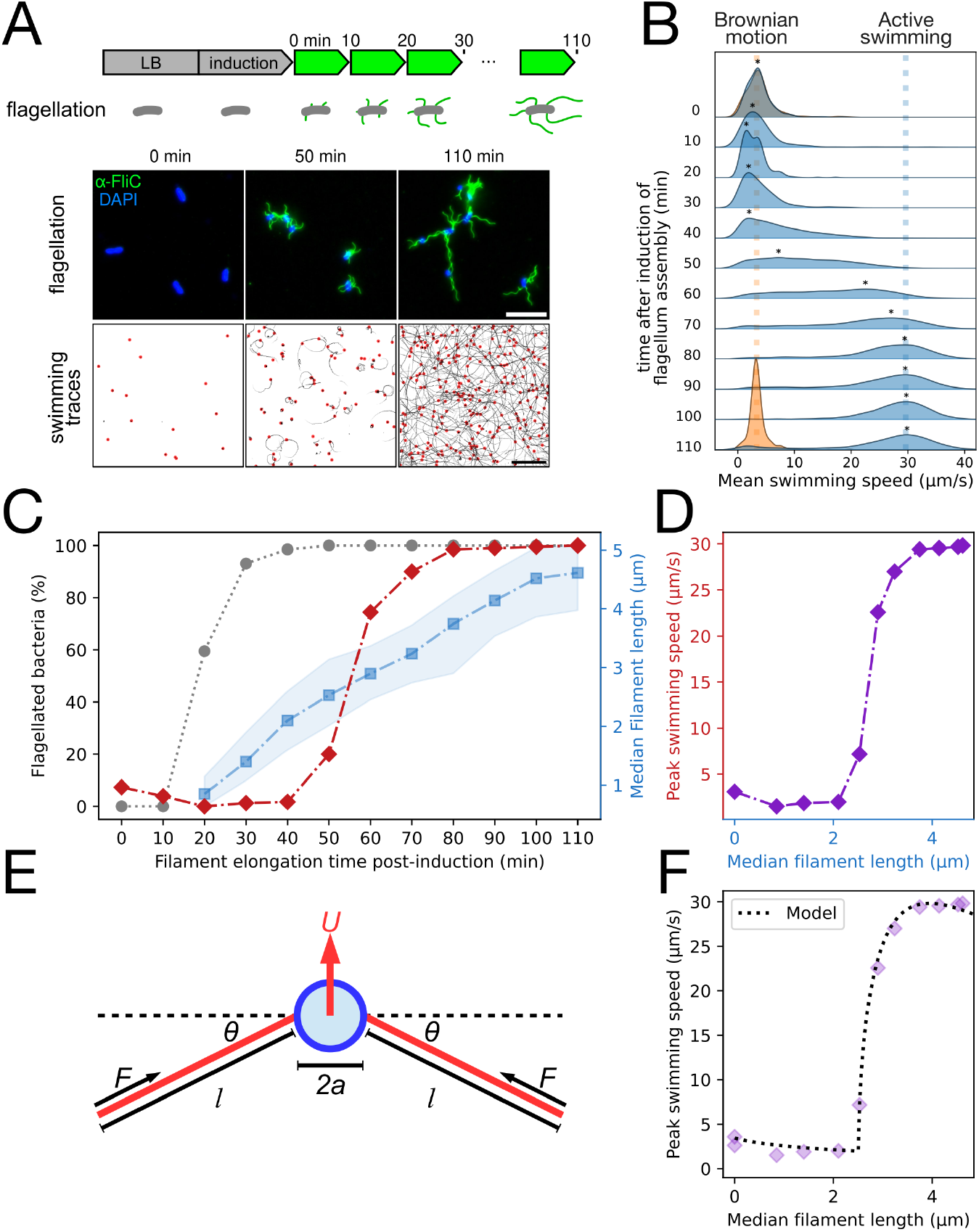
Minimal and optimal flagellar filament length to enable swimming motility. **A.** Schematic of the experimental setup and exemplary images of the immunostaining for flagellar length measurement (upper panel, scale bar = 10 µm) and swimming speed measurement and trajectories identified using iLastik and TrackMate (lower panel, scale bar = 50 µm). **B.** Density plot of mean single-cell swimming speed after induction of flagella synthesis. Blue density plots represent the WT swimming speed at the different timepoints, orange density plot represents the moving speed (only due to Brownian motion) of the non-motile Δ*motA::mudJ* strain at t = 0 min and 110 min after medium switch. The star (*) above the blue density plots indicates the most common (peak) speed. **C.** Proportion of flagellated bacteria (grey dots); filament length (blue) and peak swimming speed (red). The peak speed was measured as the swimming speed peak of the mean speed density of the tracks analyzed using kernel density estimates. **D.** Peak swimming speed relative to the median filament length indicate a threshold between 2 to 3 µm filament length in order to reach maximal swimming speed. **E.** Minimal physical model for a peritrichously flagellated bacterium, consisting of a body of radius *a* and flagella of axis lengths *l*. The flagella are active and each generates a parallel driving force *F*. Each flagellum is attached to the body by a torsional spring of spring constant *K* and deflection angle *θ* from the relaxed state. **F.** Fit of the physical model on the experimental data from panel D with N = 5 flagella.

As expected, no swimming above background Brownian diffusion was observed in a mutant unable to rotate its flagella (Δ*motA*) (Figure 1B and Figure S1). In contrast, for the wildtype (WT), reaching a flagellar filament length of 2.5 µm marked a sharp transition between no motility and the maximal swimming speed; below 2 µm, no swimming was observed among the population (Figure 1C and D). Interestingly, any subsequent increase in filament length was not providing any substantial speed increase, as observed when plotting the filament length across the population relative to the maximal swimming speed measured (Figure 1D).

To rationalize the physical origin of the experimentally observed continuous but rapid transition from no swimming for small flagellar filament to swimming for flagellar filaments beyond some critical length, we extended a previous physical model of the swimming transition for soft hooks (30). We modeled the cell as a rigid spherical body, to which are attached a number *N* of rigid, force-generating filaments (modelling the action of propelling, rotating flagellar filaments), each connected to the body via a torsional spring to approximate the dynamics of the flagellar hook (31) (Figure 1E). We resolved the force and torque balance on the model bacterium, ultimately obtaining an equation for the stable equilibrium angle *θ* between the axis of each rotary motor and its corresponding flagellar filament; a numerical solution allowed us to identify the stable angle and hence to deduce the corresponding swimming speed of the cell (see Supplementary Information). For a given set of physical parameters, the model invariably predicts the existence of a sharp transition to swimming at a critical flagellar filament length (Figure 1F). Below the critical length, the stable angle is *θ* = 0 and so no swimming occurs. In this regime the apparent swim speeds are actually the result of displacement due to Brownian motion. Once the flagellar length surpasses the critical length, *θ* sharply increases. Physically, this is when bundling of flagellar filaments is taking place, resulting in swimming of the whole cell and can be understood as an elasto-hydrodynamic instability. Beyond rationalizing the existence of an instability, our model can be used to infer the value of the hook stiffness in swimming bacteria. Many of the geometrical and dynamic parameters in the model are either known or can be determined theoretically (see Supplementary Information). However, the critical flagellar length (*l_c_*) and the flagellar propulsive force (*F*) generated by flagella can be determined by fitting the theoretical prediction against the experimental data. The results are shown in Figure 1F for *N* = 5 flagella. Our theoretical model predicts *l_c_* ≈2.51 *µ*m and *F* ≈1.10 pN. In turn, these can be used to deduce the value of the spring constant of the hooks, and consequently their bending modulus, which we find to be *EI* ≈5.2 ·10^−26^ N ·m^2^; the model can be further used to calculate the torque generated by the rotating motors and the angular rotation rate of the flagella (see Supplementary Information). We calculate a torque of 1.1 · 10^−18^ Nm, consistent with previous observations (32). Angular rotation rate varies between experiments, depending on flagella length *l* and fluid viscosity *µ*, but generally agrees with previous observations, typically a bit more than 100 Hz (32, 33). Notably, the value of *EI* we determined here is smaller than previous *in vivo* mea-surements. It has been established that the molecular motor twists the hook, causing changes in its molecular structure which have the effect of increasing its bending modulus *EI* during standard swimming (8, 31, 34). Strikingly, the value of *EI* in our data appears smaller than expected, being associated with very low levels of twist (8, 31). It is possible that the compressive force *F* acting on the hook causes molecular changes which act to reduce the bending modulus in contrast to the twisting, which we see is beneficial for triggering the elasto-hydrodynamic instability necessary for swimming but could also be useful for the purposes of initiating a tumble (31). Another possible explanation is the effects of Brownian motion on individual cells, which could allow the elasto-hydrodynamic instability to occur more easily, which would effectively be similar to a reduction in *EI*. The most notable prediction of our biomechanical model is the existence of a theoretical maximum swimming speed, which coincides with flagellar filament lengths observed experimentally (Figure 1F), suggesting that swimming speed is the parameter that the bacteria choose to optimise.

### The injection-diffusion model predicts a narrow range of *k*_on_ for optimal filament elongation

As shown above, we observed that the commonly referenced 20 µm long bacterial filament was not a prerequisite to achieve maximal swimming motility. A short filament was enough to achieve a quasi-maximal function, suggesting that the time needed to reach this threshold length is the physiologically important biological parameter. Accordingly, we next set-out to theoretically and experimentally rationalize the effect of the filament elongation rate on the onset of motility. Filament elongation is a non-linear process driven by two major forces: injection of the flagellins by the export apparatus and diffusion in the hollow filament (18). The injection rate *k*_on_ was previously predicted to be about 30 flagellin subunits per second by fitting the injection-diffusion equation to experimental data obtained by fluorescent multi-labeling of elongating flagellar filaments (18). Alternatively, it is also possible to estimate the elongation rate by measuring the length increment over a period of time. In order to rationalize the effect of *k*_on_ on the biological motility parameters, we used the injection-diffusion model to calculate (i) the theoretical time to reach the threshold length required for motility, (ii) filament length after 20 minutes of elongation, (iii) the gain in time to reach motility upon an increase (doubling) of *k*_on_, and finally (iv) the increase in *k*_on_ that would be required to reduce by few seconds the duration of assembly of a short filament (Figure 2). All four investigated parameters suggest that optimal filament elongation rate occurs for a *k*_on_ between 20 and 50. For *k*_on_ values below 20, bacteria would have an incentive to spend extra energy in order to achieve motility earlier. For *k*_on_ values above 50, the high injection force does not allow any significant improvement of the elongation kinetics and the incentive is rather to reduce the *k*_on_ to avoid wasting energy (Figure 2).

**Figure 2.**
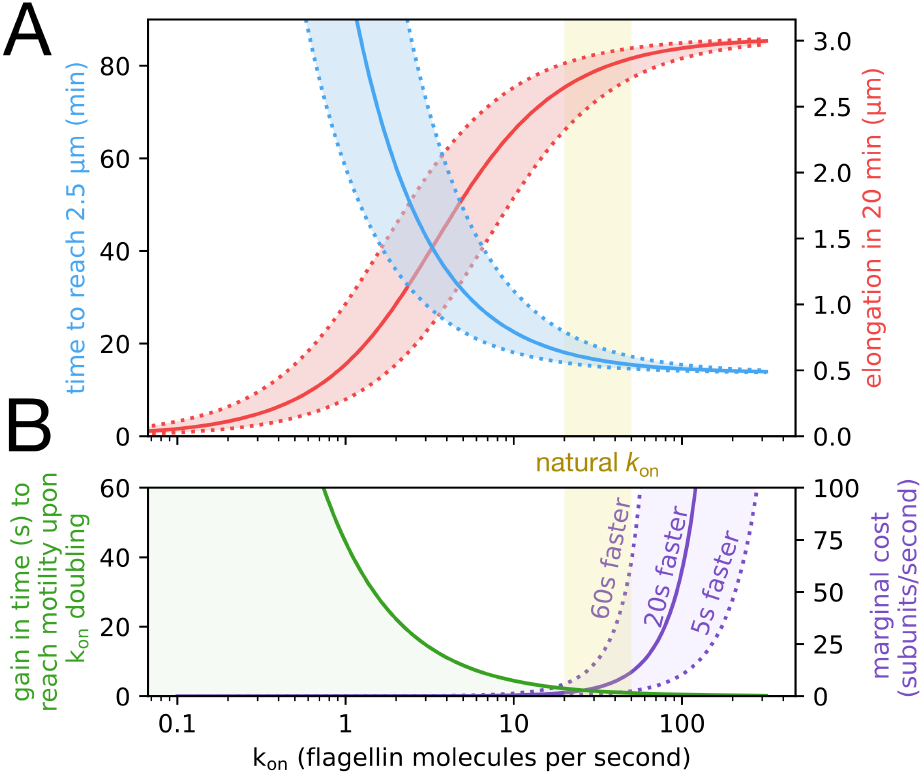
Injection by the fT3SS is a very fast process that is optimised for cellular energy conservation. **A.** The natural *k*_on_ value (yellow shaded area) is an optimum between fast elongation speed and energy conservation. Filament elongation over about one generation (20 min, in red) and the time required to reach the minimal length suitable for motility (2.5 µm) increase and decrease, respectively, with *k*_on_. Both values approach the asymptotic extremum for *k*_on_ ≈ 20–50 s^-1^, above which a plateau is reached. The dotted curves show the effect of a ±2 times fluctuation of *k*_on_ around its value. **B.** Below a few tens of flagellin subunits injected per second, any increase in injection speed significantly reduces the time to reach motility. Above *k*_on_ ≈ 50 s^-1^, the gain in time drops below a second. Accordingly, the marginal cost (extra subunits to inject per second to reach motility sooner) significantly increases above this value. This is shown here for a gain in motility onset of 5s, 20s, and 60s. NB. it takes ∼15–18 minutes to reach motility for *k*_on_ ≈ 20–50 s^-1^.

### Improving the estimation of *k*_on_ using electron microscopy to measure the filament elongation speed

The injection-diffusion model suggests that an optimal *k*_on_ exists, which reconciles fast elongation and energy-efficiency. We next aimed to experimentally determine this value. We first set out to determine the optimal temperature for filament elongation. Employing previously described stepwise filament labeling (18), we determined the growth rate of multilabeled flagellar filaments of bacteria cultivated in a range of temperatures from 25 °C to 40 °C. To facilitate filament length quantification, we developed a new method to analyse fluorescently labeled filaments using the Fiji plugin MicrobeJ (see Material and Methods section). Multilabeling analysis showed an increase in filament growth rate as the temperature increased (Figure 3A and B, Figure S2). Utilizing the mathematical equation from the injection diffusion model, we plotted the initial injection rate *k*_on_ and diffusion co-efficient D across different temperatures. Similar to the growth rate of the filaments, *k*_on_ and D increased from 25 °C to 38 °C. Above 38 °C, a decrease in both parameters can be observed, most likely caused by heat stress (Figure 3B).

**Figure 3.**
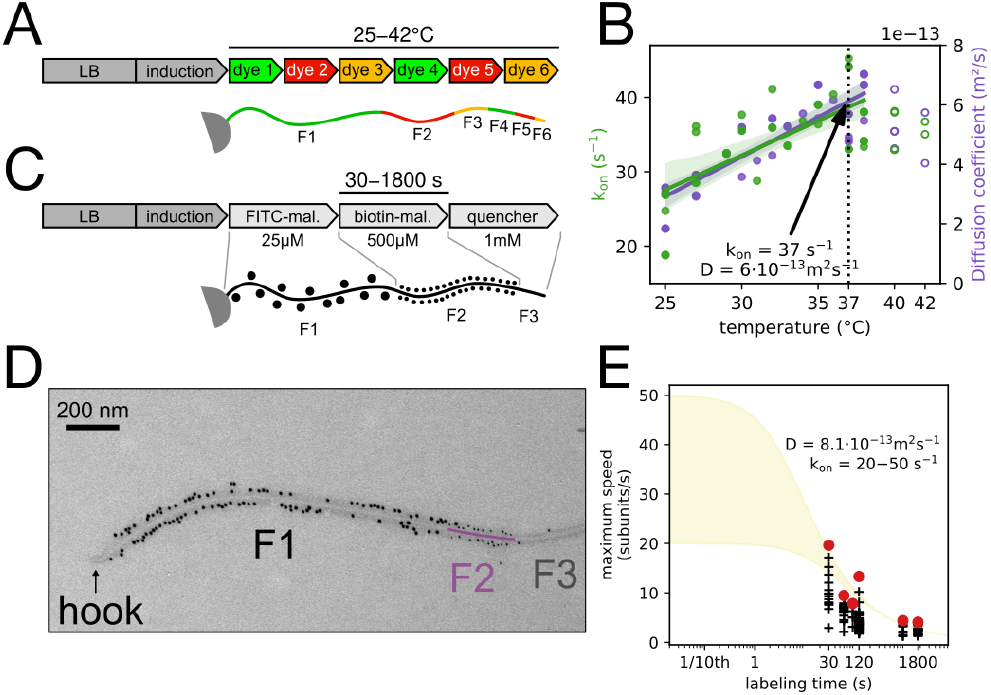
stepwise high-resolution filament labeling discloses maximal fT3SS injection rate. **A.** Multilabeling setup of flagellar filaments as previously described in (18). **B.** The injection rate of the fT3SS (*k*_on_), and the diffusion coefficient of the substrates in the channel (D) increase linearly with temperature up to 38 °C. Above this temperature, heat stress affects secretion. The arrow shows the parameters at the physiological temperature of *S. enterica* (37 °C). **C.** An improved protocol of stepwise filament labeling allows more precise measurement of filament growth over short periods of time. Use of gold beads of different sizes enables the separation between the basal fragment (F1) and the apical fragment (F2). **D.** Exemplary EM image of a flagellar filament with 60 seconds staining of the apical fragment. **E.** Experimentally measured speed of elongation of apical fragment for which the length of the basal fragment F1 is comprised between 0 and 1 µm. Decrease of the average subunit secretion rate is observed as the labeling time increases. The maximal rate recorded for each elongation time (red dots) is in agreement with the injection-diffusion model. The quadratic nature of the growth kinetics would require labeling times of less than a second to measure secretion rate close to the theoretical maximum (*k*_on_). The yellow area represents the expected flagellum length for *k*_on_ ≈ 20–50 s^-1^.

Accordingly, we measure the maximal filament growth rate at 37 °C. Given the non-linear and rapidly decreasing filament growth kinetics, the previously used stepwise filament labeling method was not suitable for determining the maximal elongation rate (*k*_on_). Moreover, the effectof biological variability and the underestimation of the measured speed is much more pronounced as one gets closer to the initial secretion events. We estimated that reducing the labeling time to less than a minute could enable us to a measure secretion rate that approaches half of the theoretical *k*_on_ value. However, this required us to improve both the spatial and temporal resolutions over previous stepwise filament labeling methods. Spatial resolution was increased by using electron microscopy (EM) instead of fluorescence microscopy. This allowed us to measure fragments with a precision of ∼ 10 nm (vs. a few hundred nanometers previously). Temporal resolution was improved by optimizing the protocol to remove any waiting steps (such as centrifugations and washes). In brief, we labeled successive fragments with increasing concentrations of two maleimide compounds, bound to either fluorescein (FITC) or biotin, to label the basal fragment (F1) and the apical fragment (F2) successively. After labeling the apical fragment, the reaction was stopped using an excess concentration of a quencher (DTT). FITC- and biotin-labeled fragments were immunostained using antibodies coupled with gold particles of different sizes (10 nm and 5 nm, respectively), observable using EM and enabling the separation between apical/basal fragments (see Material and Methods section). This approach facilitated precise labeling of filaments for durations ranging as short as 30 seconds to as long as 1800 seconds (Figure 3C and D). As we aimed to measure the subunits secretion as close as possible to the maximal elongation speed, we analyzed labeled apical fragments for which the basal fragments were in the early stage of assembly, with a length shorter than 1 µm. As predicted, decreasing labeling times revealed much higher flagellin secretion rates than previously measured (Figure 3E). We recorded a maximum *k*_on_ value of 20 flagellin molecules per second – almost 10,000 amino acids per second – which is more than 6 times faster than our previous measurements (18). We note that the measured flagellin secretion rate of 20 subunits per second is about half of the theoretical maximum, experimentally validating that secretion reaches extremely fast rates in the early stages of filament elongation. We hypothesize that the observed variability in average speed for a given elongation time is due to the cell to cell biological variability in the available pmf and cytoplasmic pools of flagellins.

### The microscopic mechanism of inserting a flagellin molecule in the fT3SS secretion channel and the energy cost for achieving an injection rate *k* _on_

To understand the biological implications of the observed rapid flagellin secretion rate, we next developed a biophysical model of flagellum growth, which correlates the energy costs of flagellin secretion with our swimming motility measurements. In this model, the injection rate *k*_on_ represents the speed at which a partially unfolded flagellin is fully inserted into the channel. This process involves two primary steps: (i) The unfolded flagellin (FliC) arrives at the entry point of the basal body; (ii) The unfolded flagellin is inserted segment-by-segment into the channel by a pmf-powered secretion system located at the basal body of the flagellum. Together the arrival and insertion timescales, *t_a_* and *t_i_*, determine to the overall injection rate, *k*_on_ = (*t_a_* + *t_i_*)^−1^. We have developed microscopic models for the mechanisms underlying both arrival and insertion steps of the injection process, which allow us to estimate the associated time scales and energy costs (full details in Supplementary Information). Our results provide insights into how the bacteria can improve their motility by spending more energy to increase the injection rate *k*_on_ and show that the experimentally observed range of *k*_on_ lies within the range of optimal efficiency.

The insertion step involves balance between the pmf, which pushes the flagellin into the channel segment-by-segment, and an entropic force that tends to push the flagellin out of the channel (Figure 4A). The later force arises because the fully extended flagellin in the channel is a high entropy state compared to outside the channel where the unfolded molecule has access to many more configurations. The insertion speed is determined by these forces along with the drag coefficient *µ*_0_ per unit length of inserted flagellin molecule. To estimate this, we use the Einstein relation, *µ_tot_* = *µ*_0_*L* = *k*_B_*T/D*, where *L* = 74*nm* is the total extended length of the flagellin, *k*_B_ is Boltzmann’s constant, *T* is temperature, and *D* = 6 · 10^−23^ *m*^2^*s*^−1^ is the diffusion constantin the channel, which is fitted by using the injection-diffusion model and the flagellar filament elongation rate measurements as described in the preceding sections. We find that the drag coefficient is very small, *µ*_0_ = (2.3 · 10^−8^*s/nm*^3^)*k*_B_*T*. For typical pmf energy scale 5–10 *k*_B_*T* and assuming the flagellin is inserted in segments of 1–2*nm*, this leads to a total insertion timescale of *t_i_* ≈ 10^−4^ *s* (see Supplementary Information). Comparing to the experimental estimates above, *k*_on_ ≈ 20–50*s*^−1^, we see that insertion is not the dominant timescale and therefore is not a useful target for improving performance. To summarize, it is important that the pmf can overcome entropic forces that tend to push a partially inserted flagellin out of the channel, but as long as the pmf is sufficiently strong the insertion will be extremely fast.

**Figure 4.**
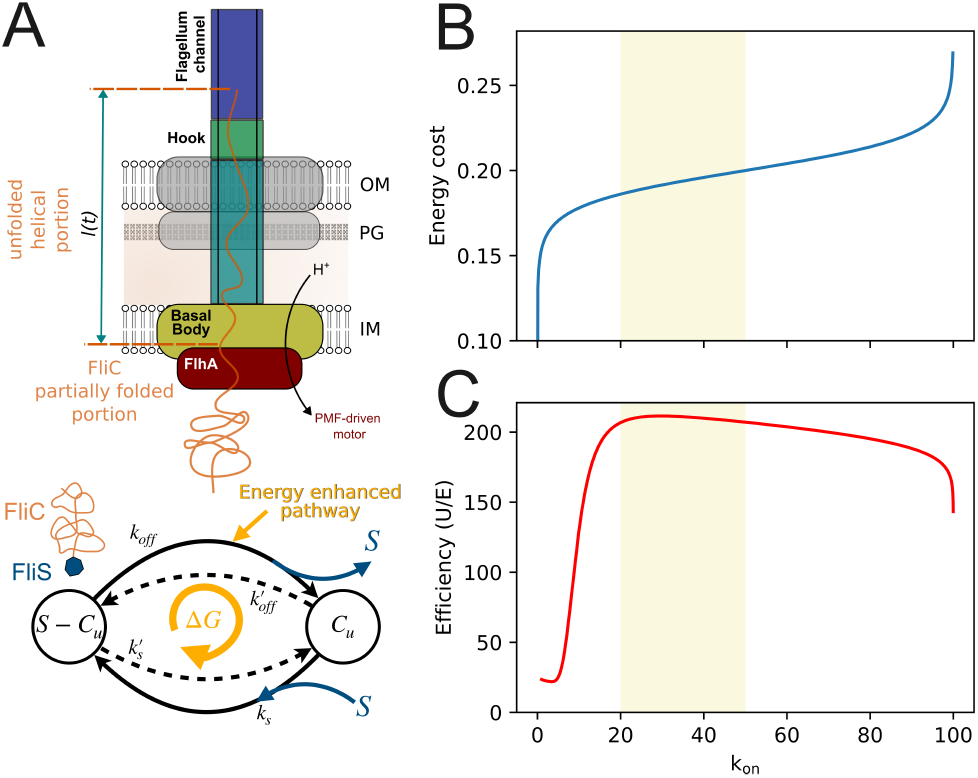
Injection by the fT3SS is a very fast process that is optimised for cellular energy conservation. **A.** The energy-rate relation for the flagellin injection process via the fT3SS. The schematic illustrates the active injection process powered by a pmf-driven motor and a minimal model for the flagellin arrival at the export gate. We consider two primary states for flagellin: partially unfolded FliC bound to FliS and free FliC available for export. Transitions between these states involve association/dissociation between chaperon and substrate in solution (with rates *k_s_* ≫ *k*_s_′) and a second pathway that strips off the chaperon via energy consuming interactions with the ATPase and/or a pmf powered mechanism. **B.** The energy cost per arrival *E*_a_ predicted by our model and **C.** efficiency *η* = *U/E* plotted as a function of the resulting injection rate *k*_on_. Here the utility function *U* is the measured average swimming speed as a function of flagella length. Increasing the injection rate requires larger energy cost that is used to increase the local concentration of FliC near the export gate. This combined with the diminishing returns for swimming speed leads to a peak efficiency within the shaded range corresponding to experimentally measured injection rates, *k*_on_ ≈ 20–50*s*^−1^.

Given the above analysis, the arrival step is therefore rate-limiting for determining the overall injection rate *k*_on_. After synthesis, flagellin binds to the chaperone FliS, which keeps the flagellin in a partially unfolded state suitable for insertion by the pmf as described above. The chaperone is stripped-off before insertion, which incurs an energy cost. This had been thought to be mediated by ATP hydrolysis by the ATPase complex, but mutants without ATPase or suppressed ATPase activity are able to export FliC (14, 15). It is therefore plausible that the pmf is at least partially responsible for chaperone removal, perhaps via interactions with the FlhA ring before the unfolded and unchaperoned flagellin reaches the export gate. Because of this uncertainty in the microscopic mechanisms, we employ a simplified but general two-state kinetic model pictured in Figure 4A. In solution, flagellin heavily favors binding to FliS with association and dissociation constants *k_s_* and *k*′_s_ respectively (*k_s_* ≫ *k*′_s_). A dissipative mechanism spends Δ*G* free energy per cycle to enhance a second reaction pathway that strips off FliS (with rate *k*_off_), leaving an unfolded flagellin molecule, which can either be exported or rebind to FliS. By spending more energy driving this chemical cycle, the bacteria can increase the local concentration of unfolded flagellin, [*C*]*_u_*, near the export gate to decrease the arrival time. Assuming the arrival is a diffusion limited process, the arrival time is *t_a_* = (4*πdD_c_*[*C*]*_u_*)^−1^, where *D_c_* is the diffusion constant for the unfolded flagellin and *d* is the interaction length scale with the pmf export complex (how close the molecule must diffuse to guarantee it is exported). Replacing [*C*]*_u_* the total intracellular concentration of FliC gives a lower bound *t_a_ > t_min_* ≈ 0.01*s*, which is only about three times smaller than the measured 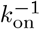. Using our kinetic model, we next compute the energy dissipated per flagellin export, *E*, and express this in terms of the resulting injection rate *k*_on_ (see Supplementary Information). As shown in Figure 4B, increasing *k*_on_ requires a greater energy cost with diminishing returns: infinite dissipation is required to push toward the limiting value 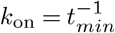. In the opposite limit, a finite, but very small, *k*_on_ can be achieved with no energy cost due to spontaneous unbinding of the chaperone. Finally, combining this energy cost with our measurements of swimming motility, we compute the efficiency, defined as *η* = *U/E*, at which the bacteria spend energy for enhancing flag-ellum function (Figure 4). For the utility function, *U*, we use the peak swimming speed as a function of filament length. Due to the combination of increasing energy cost to increase *k*_on_ and the diminishing returns in utility once the flagellar filament elongation rate is sufficiently large, the efficiency peaks around *k*_on_ = 20–50*s*^−1^; indicating that the experimentally estimated value may be tuned toward optimal efficiency.

## Discussion

Flagellum assembly in bacteria is a tightly controlled process that involves a remarkably fast secretion rate of thousands of amino acids per second through the fT3SS, a pmf-dependent mechanism that is considerably more rapid than the general protein secretion system (Sec system) which secretes only a few dozen amino acids per second and any other known pore-based protein secretion system (21, 35). The physiological importance of such a rapid protein secretion rate remained unclear. However, it appeared reasonable to speculate that flagellin secretion rate was evolutionary optimized to balance the energy costs of the secretion process and to minimize the time needed to achieve motility. Indeed, when we modeled various biologically relevant parameters fordifferent values of the flagellin secretion rate (*k*_on_), we identified a critical *k*_on_ threshold between 20 and 50 s^-1^. Beyond this threshold, increasing *k*_on_ did not further enhance elongation speed or decrease filament assembly time, but significantly increased energy costs, suggesting an evolutionary optimization of the flagellin secretion rate to reach the required minimal filament length for motility without excessive energy expenditure (*i.e.* consumption of the proton gradient and the production of flagellin). We therefore determined experimentally the minimal filament length needed for motility after induction of flagellar synthesis. Although the flagellar filament can grow to a length of up to 20 µm, we observed that a length slightly exceeding a critical value in the 2–3 µm range was sufficient to attain the maximum swimming speed (Figure 1). We note that this result is consistent with previous observations that a single helical pitch of the flagellar filament, approximately 2 µm in length, was sufficient to enable bacterial motility (28, 27). Our biophysical model, which effectively simulates the complex biomechanics of flagellar propulsion, reveals that the emergence of swimming at a specific filament length can be attributed to an elasto-hydrodynamic instability of the swimming cell with flexible flagellar hooks. This phenomenon illustrates a mechanical interplay between the hook’s elasticity, aligning the flagellar filaments with the motor axis, and the external hydrodynamic moments induced by swimming flows. The short filaments initially resist these external forces due to the stabilizing elasticity of the hooks. However, as the filaments elongate, they reach a critical length where the external forces overpower the hook’s resistance, thereby enabling swimming.

To understand the energy cost of filament growth, we developed a microscopic model for flagellin insertion into the secretion channel. Two critical steps delineate the injection process characterized by *k*_on_: the arrival of the unfolded flagellin and its segment-by-segment insertion, influenced by both pmf and entropic forces. We found that the rate-limiting step is the arrival of unfolded FliC, indicating that this step is a better target for spending energy to increase the overall injection rate. The total energy cost for each cycle of flagellin export, combined with the experimental measurements of swimming motility, allowed us to determine an optimal efficiency peak for the injection rate *k*_on_, reflecting a finely tuned balance that maximizes bacterial motility with minimal energy expenditure. This optimized behavior may reflect evolutionary adaptations that enhance bacterial fitness during swimming motility.

Within the T3SS, one notable instance of optimization is the tightly regulated secretion process that ensures the correct order of component assembly. The T3SS must secrete the rod, hook, and filament proteins in a sequential order to correctly form a functional flagellum. In other words, the T3SS is optimized to recognize and prioritize the secretion of different proteins based on the stage of flagellar assembly (36). The optimization process of the fT3SS can also be observed in the balance between the assembly speed and energy burden. Our work suggests that the fT3SS injection rate has been evolutionarily optimized to strike a balance between efficient flagellum assembly and minimal energy expenditure. This finely-tuned balance underlines the intricate evolutionary adaptations that bacteria have developed to enhance their motility and fitness in their specific environments. Our findings also raise questions about the secretion capabilities of virulence-associated type-III secretion systems. If a high injection rate is possible for the flagellar T3SS, it would be interesting to investigate whether similar rates can be achieved in the secretion systems responsible for assembling virulence factors and effectors. A previous study reported a secretion rate of 5–60 molecules per seconds of the fT3SS evolutionary-related vT3SS injectisome effector SipA in eukaryotic cells during infection, which is in range with the secretion rate of the flagellin measured during our observation. However, it remains unclear if those numbers result in the secretion of one or several vT3SS attached to the eukaryotic cell (37). Further, investigation of the observed temperature-dependent effects on secretion rate via the T3SS and the secretion kinetics of other secretion systems could yield important insights into how bacteria have evolved to optimize their secretion systems for survival in various environments and host interactions.

In conclusion, the findings presented in this study suggest that flagella assembly in bacteria has evolved to enable a swift onset of motility, crucial for survival in changing environments. The bacterial flagellum therefore represents a paradigm example of evolutionary adaptation. Its assembly requires a finely-tuned regulation, balancing assembly speed and energy expenditure, thereby showcasing the refinement process of biological optimization.

## Materials and Methods

### Strains, media and growth conditions

Strains and genotypes are listed in Table S1. *Salmonella enterica* serovar Thyphimurium LT2 was grown in LB (Lennox) (10 g/L tryptone, 5 g/L yeast extract, 5 g/L NaCl) and at 30 °C unless stated otherwise. For protocols requiring washing steps, phosphate-buffered saline (PBS) (8 g/L NaCl, 0.2 g/L KCl, 1.15 g/L Na_2_ HPO_4_, 0.2 g/L KH_2_ PO_4_, pH 7.3) was used.

### Swimming behavior and filament immunostaining

Overnight cultures were incubated in LB at 30 °C and 180 rpm. Subcultures were inoculated 1:100 in 10 mL fresh LB and cultivated accordingly. After 2.5h of growth, *flhDC* expression was synchronized by induction with AnTc, (final concentration = 100 ng/ml) followed by 30 min of incubation. Subsequently, cells were harvested at 2,500 × *g* for 5 min and resuspended in fresh AnTc free media. Directly, at 0 min post medium switch a first sample was drawn and processed for obtaining swimming behavior and filament immunostaining. Cells were placed back for incubation and new sample acquisition was performed every 10 min up to 110 min post medium switch. For each sample at the indicated timepoints, cells were diluted with LB (1:20 or at timepoints beyond 40 min post medium switch 1:40) to keep cell density below OD_600_ = 0.1. For obtaining swimming behavior of the cells, 70 µl were loaded in a flow cell and microscopy was performed using a Ti-2 Nikon inverted microscope equipped with a CFI Plan Apochromat DM 20× Ph2/0.75 objective. For each timepoint two positions were monitored for 100 frames with a time interval of 43 ms between frames. Image analysis and tracking of cells was performed using ilastik (38) and Fiji (39) equipped with TrackMate (40). Mean swimming speed was measured with TrackMate and peak swimming speed was defined as the maximum speed of a kernel density estimate from the single cell trajectories from 100 bins. The LoG detector settings were set to three microns and a threshold of 20, combined with the auto function. For the simple LAP tracker the maximal linking distance was set to two as well as the values for maximal gap-closing distance and frames. Then, a minimal displacement of three microns was determined as threshold distinguishing between movement and drift. For each timepoint in parallel cells were mounted for immunostaining as described previously (41). In brief, poly-L-lysine coated coverslips were flushed with 100 µl cells followed by fixation with 4% paraformaldehyde for 10 min. All steps were performed at room temperature. After that, cells were washed with PBS and blocked with 10% BSA for 10 min. Primary antibody (anti-FliC, rabbit, 1:1,000 in 2% BSA) was added and incubated for one hour. Subsequently, cells were washed and blocked again as described above. Secondary antibody (anti-rabbit Alexa Fluor488, diluted 1:1000 in PBS) was added and the mix was incubated for another 30 min. Finally, cells were washed twice with PBS and Fluoroshield with DAPI mounting medium (Sigma-Aldrich) was added. Fluorescence microscopy was carried out using a Ti-2 Nikon inverted microscope equipped with a CFI Plan Apochromat DM 60× Lambda oil Ph3/1.40 (Nikon) oil objective. Microscope settings were set to: 488 nm: 200 ms 2% 16 bit Z-stack every 0.3 µm range 2 µm 9 slides 405 nm: 50 ms 20% 16 bit single plane PC: 100 ms 80% DIA 16bit single plane. Image analysis (filament counting and length determination) was performed using Fiji (39) equipped with the MicrobeJ plugin using the ROI Manager for measurement of the length (42).

### Temperature variation and multilabeling of the flagellar filament

Filament multilabeling was performed as described previously (18) and as shown in Figure S3. A FliC_T237C_ cysteine replacement mutant was grown overnight in LB at 30 °C, diluted 1:100 into 10 mL fresh LB in a 100 mL flask and grown at 30 °C for 1.5 h until reaching early exponential phase. P*tetA-flhDC* promoter was induced by addition of AnTc (100 ng/mL) for 30 min. Afterwards, cells were collected by centrifugation for 3 min at 9,000 × *g*, resuspended in 10 mL fresh LB. Aliquots of the culture in 1.5 mL tubes were then incubated for 30 min at the corresponding temperature in a thermomixer with low agitation (300 rpm). Labeling with maleimide dyes up to 6 filaments fragment was performed successively as described previously (18). After the final labeling period, cells were resuspended in PBS and 100 µL were applied to a custom-made flow cell coated with poly-L-lysin. Cells were fixed by addition of 4% PFA for 10 min, followed by a washing step with PBS. Fluoroshield with DAPI mounting medium (Sigma-Aldrich) was added and the cells were observed by fluorescent microscopy using a Zeiss Axio Observer Z1 microscope at 100× magnification. A Z-stack was applied for every image to ensure observation of the whole filament. Fluorescence images were processed using Fiji (39) and a custom Fiji macro to fuse the Z-stack.

### Filament multilabeling analysis

Sequentially labeled flagella were detected and analyzed using MicrobeJ 5.13n (42) a plugin for ImageJ (39). Briefly, rough estimations of medial axes of flagella were manually drawn on a maximum intensity channel-projection image using the segmented-line selection tool of ImageJ. The resulting selections were then stored in the ROI Manager and imported in MicrobeJ using the Filament detection mode. The medial axis of each filament was then refined using a transversal local maximum neighbor algorithm and used to compute the geometrical and topological properties of the filament, such as its length, width, sinuosity, curvature, and angularity. The analysis of the fluorescence intensities along the medial axis of each filament was subsequently performed using the Feature option called *Multisection* in MicrobeJ. Briefly, the fluorescent profiles were extracted along the medial axis of the filament for each specified channel and filtered using a window moving average filter. To accommodate global variations of fluorescence intensity between channels, fluorescent profiles were weighted using specific factors for each channel. Sections were defined as the brightest sections of each channel. The relative localization, the length and the relative order along the corresponding filament were determined for each section (Figure S3).

### High-resolution EM stepwise filament labeling

The previously described protocol (18) was modified as follows. Overnight cultures were incubated in LB at 37 °C and 180 rpm. Subcultures were inoculated 1:100 in 10 mL fresh LB and cultivated accordingly. Cells were grown until an OD600 of 0.6. Expression of *flhDC* was synchronized by induction with AnTc (final concentration = 100 ng/ml) followed by 30 min of incubation. Subsequently, cells were harvested at 2,500 x g for 3 min and resuspended in fresh AnTc free media. To allow completion of hook-basal body assembly, cells were incubated for further 30 minutes at 37 °C. The first and second fragments were labeled by successive addition of 25 µM Fluorescein-5-Maleimide (Thermo Fisher Scientific) and 500 µM Maleimide-PEG2-Biotin (Thermo Fisher Scientific), respectively. Labeling reactions were performed at 37 °C and mild agitation. At the end of the second fragment labeling, the reaction was stopped by addition of 1 mM of DTT. The step increase in concentrations was sufficient to precisely determine the limits of the measured fragments. The flagella were then detached from the cell by 10 passes through a 23-gauge needle and the cells were removed by 10 min centrifugation at 3,000 × g. The labeled filaments were applied to carbon and pioloform-film-coated gold grids. FITC- and biotin-labeled fragments were immunostained using anti-fluorescin (Aurion) and anti-biotin (British Biocell) antibodies coupled to 10 nm and 5 nm gold particles, respectively. Of note, due to slight chemical differences in fluorescein and FITC structures, the immunostaining efficiency for the first fragment and thus the gold coverage was not fully optimal, which had no impact on the precision of the second fragment measurement. The samples were acquired on a Zeiss LEO 906 operated at 100 kV and images were recorded with a digital camera. Only intact filaments with both the hook attached and three fragments (10 nm labeled, 5 nm labeled, and unlabeled) were used for the quantification. Fragment length was measured using Fiji (39) and the NeuronJ plugin (43).

## ACKNOWLEDGEMENTS

We thank the Erhardt and Charpentier laboratories for continuous help and support and Heidi Landmesser for expert technical assistance. This work was supported in part by a project from the European Research Council (ERC) under the European Union’s Horizon 2020 research and innovation program (grant agreement n° 864971) (to M.E.), and by a fellowship from the Alexander von Humboldt foundation (to T.T.R.). The work of D.H. and Y.T. is supported in part by a NIH grant (R35GM131734) to Y.T. The funders had no role in study design, data collection and analysis, decision to publish, or preparation of the manuscript.

## AUTHOR CONTRIBUTIONS

T.T.R. and M.E. conceived the project and designed the study; M.H., S.F., P.F.P., C.G. and T.T.R. performed the experiments; M.H, S.F., P.F.P., D.H., J.S., E.L., Y.T., M.E. and T.T.R. analyzed and interpreted the data; A.D. developed the MicrobeJ plugin for filament multi-labeling analysis; M.H., S.F., D.H., J.S., Y.T., T.T.R. and M.E. wrote the paper; E.C., M.E. and T.T.R. contributed funding and resources. All authors commented and approved the manuscript.

## Supplementary Information

**Figure S1.**
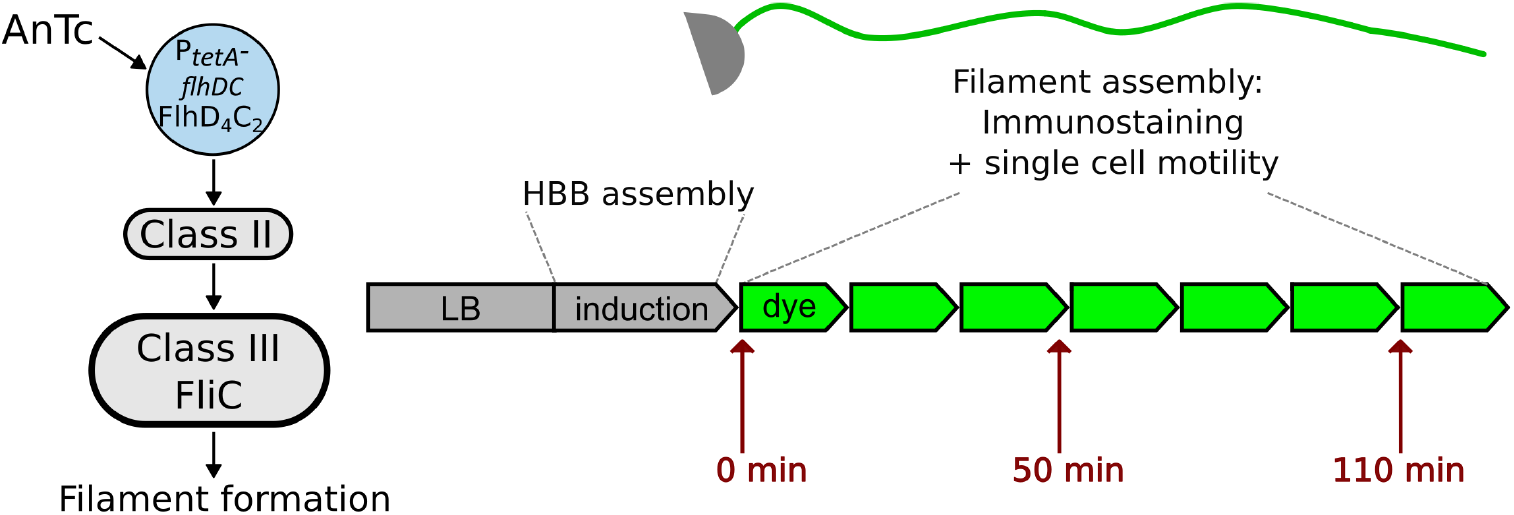
Experimental setup of the fT3SS assembly synchronization in the cell population. (**A**) After growth in LB medium, the master regulator *flhDC* expression was induced by addition of AnTc for 30 min, synchronizing expression of *flhDC* followed by class II and class III and the assembly of hook basal body (HBB) and filaments in the bacterial population. After induction, cells were resuspended in LB-medium free of AnTc inducer. Samples were probed every 10 min from the culture, simultaneously filament stained (immunostaining, rabbit α-FliC primary antibody, mouse α-rabbit secondary antibody coupled to Alexa488) and monitored for swimming speed (see Material and Methods).

**Figure S2.**
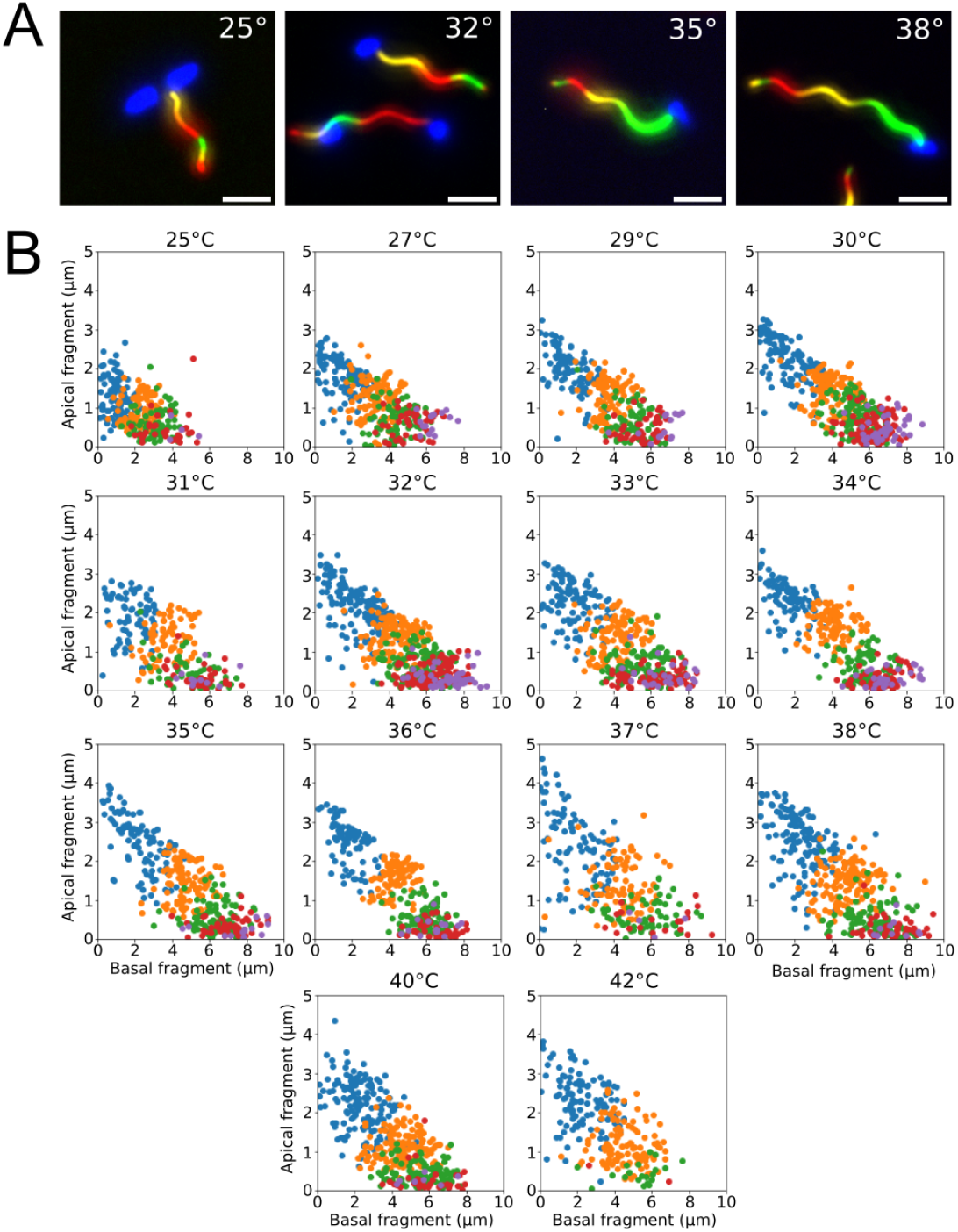
Analysis of multilabeled filaments at different temperature. (**A**) Exemplary microscopy pictures of multilabeling experiment, as previously described in (1) (scale bar = 2 µm). (**B**) Exemplary graphs of multilabeled filaments at the different temperatures. Analysis was performed using MicrobeJ plugin as described in Material and Method. At least 100 filaments were analysed per experiment. Values were represented using a custom Python script.

**Figure S3.**
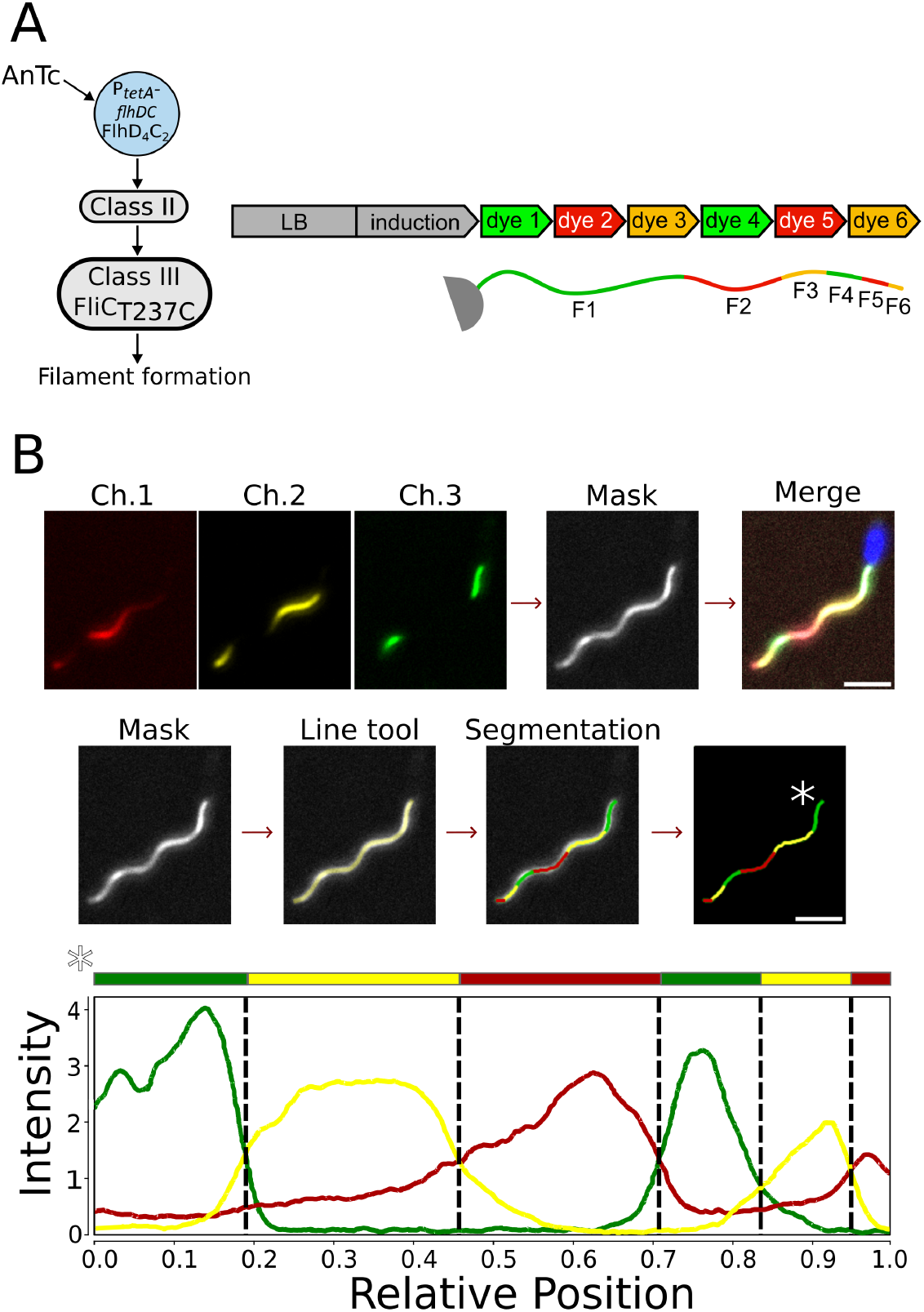
MicrobeJ analysis of the multilabeled flagellar filament. (**A**) Experimental setup shown as described in the Material and Methods and in (2). (**B**) The multi-labelled flagellar filaments were imaged successively on the red (Ch. 1), the yellow (Ch. 2), and the green (Ch. 3) channels. To facilitate the detection of the whole filaments, the channels were combined into a maximum intensity channel-projection image (Mask) using the “Combine channels” tool of MicrobeJ. The medial axis of each filament was manually drawn on the mask using the segmented line tool and then refined using a transversal local maximum neighbor algorithm. Alternatively, the blue channel was used to detect the cell attached to the filament and to define which end of the filament is connected to the cell body (*\**). The oriented fluorescent profiles were then extracted along the medial axis of the filament for each specified channel and used to segment the filament into sections (bottom graph). Sections were defined as the brightest sections of each channel. The relative localization and the length were determined for each section and represented on the filament overlay using their corresponding color.

**Table S1.** List of strains used in this study.

| Strain | Genotype | Source |
| --- | --- | --- |
| EM4869 | Δhin-5717::FRT PtetAflhDC5451::Tn10dTc[del-25] | Lab collection |
| EM11996 | Δhin-5717::FRT PtetAflhDC5451::Tn10dTc[del-25] ΔmotA5461::mudJ | This study |
| EM2046 | Δhin-5717::FRT PtetAflhDC5451::Tn10dTc[del-25] fliC6500 (T237C) | Lab collection |

### Model: Elastohydrodynamic instability of a peritrichous swimmer

#### Governing equations

A global force balance on the system gives

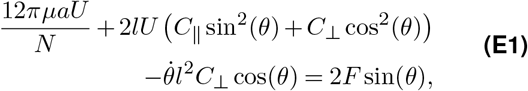

whilst a torque balance on each flagellum gives

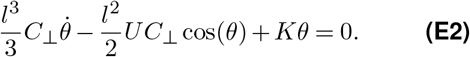

These equations can be combined for an evolution equations for *θ*. By setting 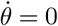 we obtain the equilibrium equation which gives the value of *θ* for given flagellar length *l*:

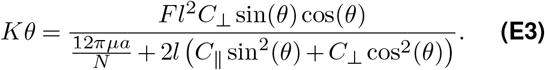

Clearly, *θ* = 0 is always an equilibrium angle. However, defining a critical length *l_c_* implicitly as

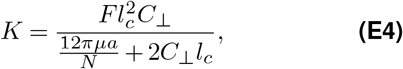

the evolution equation can be perturbed about *θ* = 0 to show that *θ* = 0 is stable if and only if *l < l_c_*. When *l > l_c_*, a new stable equilibrium at *θ >* 0 is created. Whether *l < l_c_* or *l > l_c_*, assuming the system has reached its stable equilibrium, the swimming speed can then be calculated as

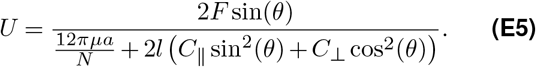

### Physical parameters

Before we can solve the governing equations, we must identify appropriate values for various physical parameters, and justify certain assumptions.

The helical pitch angle of the flagella is approximately 30^°^ (3, 4), so for an axial length *l*, the contour length is given by 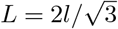. This provides appropriate definitions for the Resistive Force Theory drag coefficients

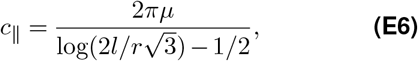

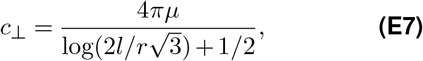

where *r* = 12 nm (4) is the contour radius and *µ* is the dynamic viscosity of the fluid. These serve as expressions for the fluid force per unit length exerted on a slender filament of length 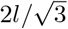 travelling at unit speed. However, the helical geometry of the real flagella complicates this. Integrating for the fluid force along the length of the contour gives total fluid forces

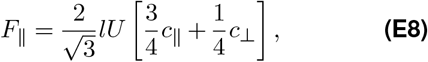

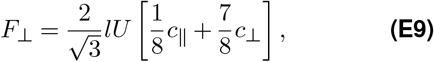

for motion parallel and perpendicular to the axis of the flagellum, respectively. This produces effective drag coefficients

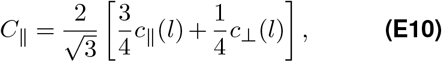

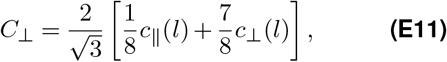

which, for clarity, give rise to drag forces acting on the flagella of axis length *l* travelling at speed *U* of

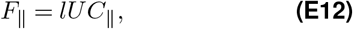

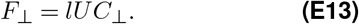

It should be noted that *C*_⊥_ and *C*_∥_ are technically functions of the axis length *l*. However, we find that over the fairly small range of values of *l* that we consider here, we can, to good approximation, set these coefficients to be constant by evaluating at, say, *l* = 3.5 *µ*m. This gives

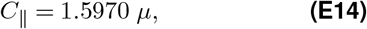

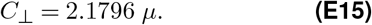

This is a fairly important result, deviating significantly from the classical result *c*_⊥_ ≈ 2*c*_∥_.

We select the radius of the body to be such that it experiences the same drag as a prolate spheroid of semi-axes 0.5 *µ*m and 1.25 *µ*m (approximating the true geometry of *S. enterica*) travelling along its long axis. The resistance/mobility matrix for such a spheroid is easily calculated (5), giving an appropriate body radius *a* = 0.65252 *µ*m. Note this slightly alters the diffusive properties of the swimmer, though only by a few percent.

Diffusion of a sphere is trivial to calculate, as a sphere in Stokes flow experiences a drag of 6*πµaU*. Using our radius *a* = 0.65252 *µ*m and an approximate temperature of *T* = 300 *K*, as well as noting a time step of *t* = 0.0435 *s* in between observation over which speed is calculated, we determine a diffusive speed of

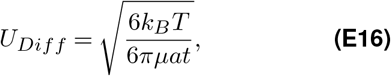

where *k_B_* is the Boltzmann constant. This diffusive speed will ultimately produce excellent agreement with the data for the initial time step. However, it is significantly more than the speeds oberserved for later precritical speeds. To correct for this, we can include the effects of the flagella. Let us suppose for simplicity (which should nonetheless provide a decent approximation of reality) that there are 6 flagella of length *l* protruding from the spherical body with uniform distribution, akin to the six faces of a die. Therefore, the swimmer remains isotropic, but for any given swimming direction, its drag has increased by an amount

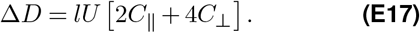

However, we wish to consider a more general number of flagella *N* (5 in our model) and so we will approximate the additional drag per unit speed as

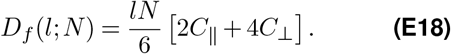

This increases the drag on the swimmer, thereby reducing its diffusive speed to

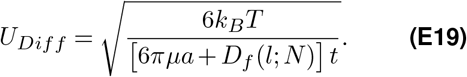

This decreases as *l* increases and gives better agreement with the data. Note that this estimate neglects hydrodynamic interactions between the flagella and the cell body, which is not expected to have a large effect. We continue to use the above expression post-criticality, even if swimming is seen to dominate diffusion throughout.

The bacteria swim in room temperature LB medium. The dynamic viscosity of LB medium is 1.83 cP when at a temperature of 37 ^°^C (6). If we assume a similar relationship between viscosity and temperature for the LB medium as there is with water, then accounting for this gives a dynamic viscosity of approximately 2.61 cP at room temperature. In SI units, this gives us an estimate for the dynamic viscosity *µ* ≈ 0.00261 Pa · s.

Applying Resistive Force Theory allows us to calculate consistent expressions for the driving force generated by the flagella in terms of either the angular rotation Ω and axis length *l*, or the motor torque *T*.

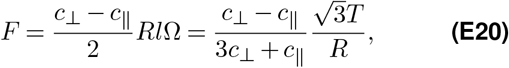

where *R* ≈ 0.2 *µ*m is the helix radius. Note we use the standard RFT coefficients, *c*_⊥_ and *c*_∥_, not the modifiedones, *C*_⊥_ and *C*_∥_. The important consequence of this is that, assuming the molecular motors generate constant torques, the force generated by the flagella are constant (within particular assumptions required for RFT to be accurate, which are satisfied post-criticality) whilst the angular rotation rate decreases as the flagella grow. This also allows us to calculate *T* in terms of *F* for comparison with pre-existing literature. The actual value of *F* is determined by numerically matching the theory to the data to minimise the square errors.The value of the bending modulus *EI* varies greatly as different values of motor torque cause twisting of the hook, reconfiguring its molecular structure and causing stiffening as the torque increases (7, 8). We identify the critical length *l_c_* in the same way as the force - by matching the theory to the data to minimise square errors. It is easy to do matching of *l_c_* and *F* simultaneously as the value of *l_c_* affects the value of *l* at which transition occurs, whilst *F* represents the size of the speeds achieved. This then allows us to calculate the torsional spring constant *K* and we then approximate the hook bending modulus by *EI* ≈*Kl_h_* where *l_h_* ≈55 nm is the length of the hook (9).

#### Final Model

When *l < l_c_*, the stable angle is *θ* = 0 and the corresponding swimming speed is *U_swim_* = 0. For *l > l_c_*, the stable angle is given by the non-zero solution to the equation

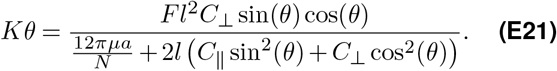

The swimming speed that results is then

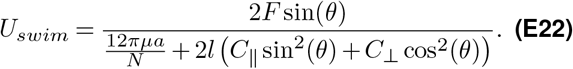

We must incorporate into this the diffusive speed

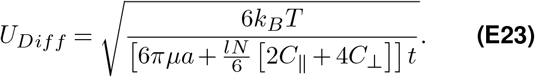

Finally, to compare with experimental data, which measured the swimming speed in a single plane, we must multiple by 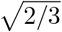 to account for the reduced degrees of freedom. We obtain the final speed

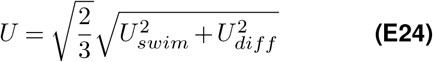

### Model: Arrival and insertion timescales and the associated energy cost

As described in the main text, the injection of flagellin into the flagella channel involves two timescales: arrival *t_a_* (how long it takes a partially unfolded FliC to arrive at the flagella base) and insertion *t_i_* (how long it takes for the pmf-powered complex to fully insert the FliC). These combine to set the overall on rate *k_on_* = (*t_a_* + *t_i_*)^−1^ for FliC. Energy is consumed in both the arrival and insertion process. Association with the chaperon FliS puts FliC in a partially unfolded state suitable for insertion; energy must be dissipated when the chaperon is removed before it arrives at the flagella base. The insertion complex then must overcome the entropic difference between the partially unfolded and nearly straight configurations. We will estimate each of the time scales *t_a_* and *t_i_* and their relation to the energy input for these functions.

#### Insertion time and energy cost

We model the insertion process using force balance between the drag force, driving force, and resistant entropic force. The former arises due to friction coefficient *µ*_0_*l* which is proportional to the current length of insertion *l*. The pmf driving force pushes the FliC into the channel with constant force *f_d_*, while an entropic force *f_r_* tends to push FliC out of the channel (where the protein has access to more configurations). Writing the driving force *f_d_* = *G*_0_*/δl* in terms of the energy *G*_0_ expended in one pmf cycle and the insertion distance *δl* per cycle, we have the following force balance equation,

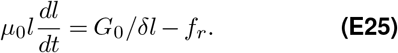

Solving for *l*(*t*), we obtain

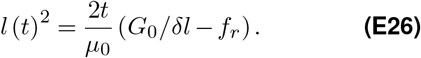

The total insertion time is defined by *l*(*t_i_*) = *L* with *L* = 74 nm being the length of the entire extended flagellin protein. Solving for *t_i_*, we find

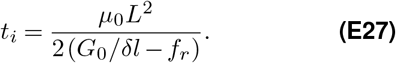

Finally, we express the total energy for insertion *E_i_* = (*L/δl*)*G*_0_ in terms of *k_on_* = (*t_a_* + *t_i_*)^−1^, treating *t_a_* as a constant (since *t_a_* is independent of the insertion energy cost),

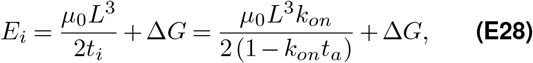

where Δ*G* = *f_r_L* is the total entropy difference between the free floating and fully inserted FliC. Notice that *t_a_* sets an upper bound max (*k_on_*) = (*t_a_*)^−1^ that is only exactly achieved with infinite dissipation by the pmf, *G*_0_ → ∞ (which leads to infinitely fast insertion, *t_i_* → 0).

To proceed, we estimate the drag constant from the measured diffusion coefficient for flagellin in the flagella channel: *D* = 0.6 *µ*m^2^/s, so that *µ*_0_ = *k_B_T/*(*DL*) = (2.3 × 10^−8^ s*/*nm^3^) × *k_B_T*. Because this drag coefficient is very small, the insertion process is very fast. The pmf energy per cycle is roughly ∼ 6*k_B_T* and we estimate the entropic energy cost for inserting by *δl* ≈ 1 − 2 nm to be around ∼ 2 − 4*k_B_T* (10). Thus, the denominator in Eq. (E27) is 𝒪 (*k_B_T*) and the resulting inser-tion time *t* ∼ 𝒪 (10^−4^ s). Compared to the measured 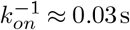, *k*^−1^ 0.03 s, we see that insertion is not the dominant timescale; spending more energy to decrease *t_i_* has negligible impact on the overall insertion rate *k_on_*. It is important that the pmf is strong enough to overcome the entropic force *f_r_*, but once this is the case, the insertion step will be extremely fast.

It is also worth noting that the excess energy cost for this fast insertion is relatively small: *E_i_/*Δ*G* = 1 + *µ*_0_*L*^3^*/* (2*t_i_*Δ*G*) ≈ 1.25 for Δ*G* = 200*k_B_T* and *t_i_* = 10^−4^ s. The cell only spends ∼ 25% more energy than if it executed the insertion adiabatically (infinitely slowly with minimal energy cost).

#### Arrival time and energy cost

From the analysis in the preceding section, we have shown that the insertion rate *k_on_* is limited by the arrival of FliC at the insertion complex. We assume the arrival time *t_a_* is diffusion limited and that once the FliC is within a distance *d* of the insertion complex (the “capture radius”) it binds with probability 1 and is quickly inserted into the flagella channel as described above. The mean first passage time for a partially unfolded FliC to diffuse to the injection site is then,

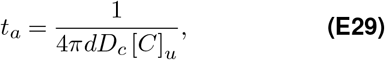

where *D_c_* is the partial unfolded FliC diffusion constant in fluid (outside the flagella channel), and [*C*]_u_ is the concentration of partially unfolded FliC. Note that [*C*]_u_ is specifically the local concentration near the flagella base, where the binding to the insertion complex occurs. The concentration need not be uniform throughout the cell if the chaperon FliS is preferentially removed near the flagella base (either by the ATPase or some pmf-powered energy consuming mechanism). Increasing the dissipation rate can drive the system to a higher concentration of partially unfolded FliC, thereby reducing the arrival time *t_a_* via Eq. (E29). If the concentration were uniform, the minimal arrival time is simply set by the total FliC concentration [*C*]_0_, *t*_min_ = (4*πdD_c_* [*C*]_0_)^−1^. On the other hand, if the ATPase activity is localized near the flagella base, we have 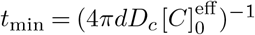, where 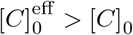 is some effective maximal concentration due to the localization of the ATPase/pmf activity. With this strategy, the cell can increase the local concentration above cell’s average concentration [*C*]_0_.

To model the energetics, we will use a simple two-state model (see main text, Fig. 4) for the populations of the chaperoned flagellin FliC-FliS and the free partially unfolded flagellin FliC, with concentrations [*C*]*_s_* and [*C*]*_u_* respectively with 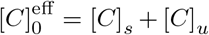. There are two transitions between these states: (1) chaperon removal with rate *k*′_off_ (with reverse rate *k*′_off_) likely involving the ATPase or a pmf powered mechanism (e.g. interactions with the FlhA ring before the FliC reaches the export gate) and (2) FliC that is not injected can rebind to FliS, with rate *k_s_* (and reverse rate *k*′_s_). There are clearly sub-steps within this cycle, for example binding to the ATPase/FlhA or the FliC folding before binding to a new chaperon. However, because little is known about these mechanisms (and the associated reaction rates), we opt for the minimal coarse-grained model. Each of these reactions are highly irreversible so the FliC tend to follow the cycle clockwise between the two states.

The model has steady state concentrations,

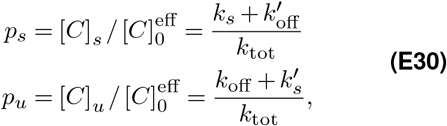

where *k*_tot_ = *k_s_* + *k*′_s_ + *k*_off_ + *k*′_off_. The dissipation rate of this system is given by,

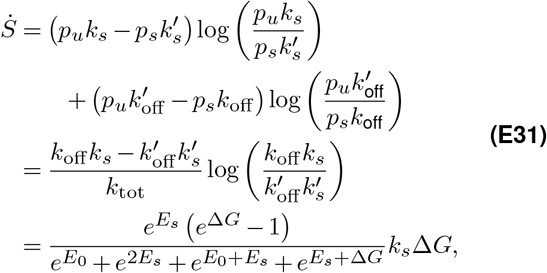

where in the final line we introduce 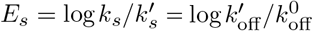 and 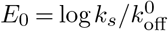. Here 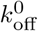 is the “bare” rate for the chaperon removal pathway with no energy input. This rate is enhanced by the dissipative mechanism 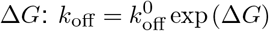. All energy scales are expressed in units of *k_B_T*. The dissipation per flagellin ejection is then equal to 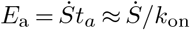. We can similarly write the arrival time (and hence 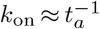) in terms of the cycle dissipation Δ*G* by combining *t_a_* = *t*_min_*/p_u_* and Eq. (E30),

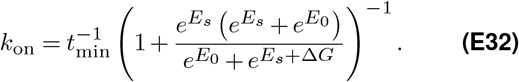

Finally, eliminating Δ*G*, we express the energy cost per flagellin ejected directly in terms of the resulting injection rate *k*_on_,

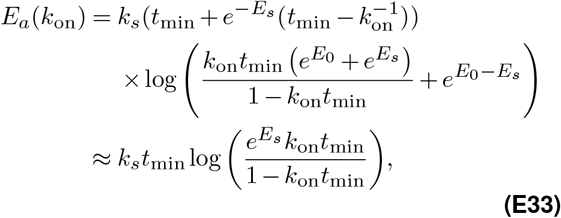

where in the final line we have taken the large *E_s_* limit to make the approximation. Notice that the energy cost diverges logarithmically as 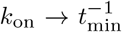, it takes infinite energy to increase *k*_on_ toward this limiting value. Conversely, the energy cost vanishes at 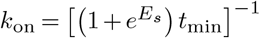. This injection rate is achieved just by spontaneous dissociation of the FliS chaperone from FliC but is extremely small because this is a rare event, *E_s_* ≫ 1.

A plot of the energy is shown in the main text (Figure 4). Here we used the following parameters: *E_s_* = 20*k_B_T*, *E*_0_ = 10*k_B_T*, and *t*_min_ = 0.01 s. The minimum time scale is roughly estimated using *D_c_* = 10 *µ*m^2^ */*s, *d* = 1 nm, and [*C*]_0_ = 1000*/µ*m^3^ leading to *t*_min_ = 0.008 s. Fig. 4 in the main text also shows the efficiency *η* = *U/E_a_*, where *U* is the measured swimming speed as a function of flagella length. To express swimming speed in terms of *k_on_* we use the length after 30 mins of growth predicted by the injection-diffusion model fit to our measurements of flagella growth. The peak efficiency falls in the measured range *k*_on_ = 20 − 50 s^−1^ and the quantitative shape of this curve is quite robust in the *E_s_* ≫1 regime. For smaller *E_s_*, the peak tends to move toward slightly smaller *k*_on_.

